# Beyond Rate: Hippocampal Phase Codes for Space and Memory

**DOI:** 10.64898/2026.10.13.758895

**Authors:** Luca D. Kolibius, Molly S. Hermiller, Mohammad F. Khazali, Stephanie Gollwitzer, Katrin Walther, Rüdiger Hopfengärtner, Hajo Hamer, Marije ter Wal, Mircea van der Plas, Simon Hanslmayr, Howard Bowman, Joshua Jacobs

**Affiliations:** Department of Neurology, University of Chicago, Chicago, IL, USA; Department of Psychology, Florida State University, Tallahassee, FL, USA; Epilepsy Center, Medical Center – University of Freiburg, Faculty of Medicine, University of Freiburg, Freiburg, Germany; Epilepsy Center, Department of Neurology, University Hospital Erlangen, Friedrich-Alexander-Universität Erlangen-Nürnberg, Erlangen, Germany; Centre for Human Brain Health, School of Psychology, University of Birmingham, Birmingham, UK; School of Psychology and Neuroscience and Centre for Neurotechnology, University of Glasgow, Glasgow, UK; School of Psychology, University of Birmingham, Birmingham, UK; Neuroscience Institute, University of Chicago, Chicago, IL, USA

## Abstract

Place cells and concept neurons are defined by a rise in firing rate at a particular location or for a particular person or object. Neurons without such a rise remain unlabeled, although they keep firing during the task. Such neurons could still carry information in the timing of their spikes. Hippocampal theta provides a reference for this timing. We therefore asked whether the theta phase of a neuron’s spikes depends on location or on the viewed image, even when its firing rate does not. We analysed hippocampal neurons in humans navigating a virtual city, rats running on a linear track, and humans viewing images of animals, faces, and places. In both navigation datasets, more neurons than expected by chance fired at a consistent theta phase at particular locations. This phase tuning remained after we excluded neurons whose firing rate depended on location. In human navigation, the spike phases of these remaining neurons improved location decoding beyond the firing rates of all neurons. More phase-tuned neurons than expected by chance also fired at progressively earlier theta phases as participants passed each neuron’s preferred location. This pattern resembles phase precession, which classically occurs as an animal crosses the firing-rate field of a place cell. The neurons that showed it had no firing-rate field, so their phase shift was anchored to the location defined by phase tuning alone. During image viewing, more neurons than expected by chance changed the theta phase of their spikes after a particular image appeared. This effect remained after we removed, for each neuron, the images that changed its firing rate. Spike phases alone also predicted the image category above chance. Theta-relative spike timing therefore carries information about location and images that firing rate alone misses. We propose that weak input changes when a neuron fires within the theta cycle, whereas strong input also raises its firing rate.

## Introduction

Canonical place cells increase their firing rate at particular locations (O’Keefe and Dostrovsky, 1971). Concept neurons increase their firing rate for particular people or objects (Fried et al., 1997; Kreiman et al., 2000), even when their appearance changes (Quian Quiroga et al., 2005). These labels made spatial and semantic representations visible at single-cell resolution, but both definitions rely on changes in firing rate. Neurons without such a rate increase remain unlabeled. They still fire during the task, but their activity is often treated as background noise. Yet neurons that are not place cells also encode position and contribute to population decoding of position in mouse dentate gyrus and CA1 (Stefanini et al., 2020). Moreover, action potentials are metabolically costly (Attwell and Laughlin, 2001), so this activity may be informative. A neuron could signal not only through how many spikes it fires but also through when it fires them.

A temporal code requires a reference against which spike timing can be measured. In the hippocampus, theta provides such a reference. Human hippocampal neurons lock their spikes to particular phases of ongoing oscillations, including theta (Jacobs et al., 2007), and stronger spike–theta coordination can predict later memory (Rutishauser et al., 2010). However, general theta locking does not establish that phase carries information about a particular location or concept: it asks whether spikes favor part of the theta cycle after activity is pooled across conditions. We instead ask whether theta-relative spike timing depends on the current location or viewed image. By testing phase structure separately across spatial locations or images, we ask whether theta timing carries specific spatial or semantic information.

We propose that spatial and semantic tuning strength varies gradually. At the strong end, a preferred location or concept increases firing enough to define a place cell or concept neuron. With weaker tuning, the same preference may be insufficient to alter the average firing rate. Because neuronal excitability varies across the theta cycle, weak input could still alter when spikes occur. A preferred location or concept could therefore produce location- or concept-specific phase structure without the rate increase required for a canonical label. Phase may thus reveal tuning in neurons that appear untuned when analysed by firing rate alone. We call a neuron a phase place cell when its spikes concentrate at a consistent theta phase at a particular location and a rate place cell when its firing rate depends on location. Phase place tuning differs from phase precession, in which spike phase advances systematically as an animal crosses the firing field of a rate place cell (O’Keefe and Recce, 1993). A neuron can be both a phase place cell and a rate place cell.

Classical phase precession established that canonical rate place cells also carry a temporal component. As an animal crosses a place field, spikes advance systematically through the theta cycle (O’Keefe and Recce, 1993). Accordingly, theta phase can improve position decoding from ensembles of rate-defined place cells, with rate and phase carrying complementary spatial information (Jensen and Lisman, 2000; Huxter et al., 2003). In human hippocampal and entorhinal neurons, phase precession was likewise identified within rate-defined fields (Qasim et al., 2021). These studies therefore show that spike timing adds information within neurons already identified by spatial firing rate. They do not test whether theta phase carries spatial information in neurons without a rate place label. We address this gap by testing location-specific phase consistency in those neurons and then asking whether their phase information improves decoding beyond firing rates from the full population.

Studies beyond classical place fields address only parts of this gap. During human navigation, spike phase distinguished prospective goals, but whether theta-relative spike timing encoded current location was not tested (Watrous et al., 2018). In lateral septum, spike phase carried position in neurons whose firing rates were spatially uninformative, but these recordings were downstream of the hippocampus (Tingley and Buzsáki, 2018). Sequence studies and phase-precession work in humans and bats show that phase organization can span nonspatial content and theta that is irregular or not continuously oscillating (Eliav et al., 2018; Qasim et al., 2021; Reddy et al., 2021; Zheng et al., 2024), although phase of firing did not reflect item order in a human sequence-memory task (Liebe et al., 2025). It therefore remains unknown whether hippocampal neurons across species include phase place cells, whether phase place cells occur without detectable rate tuning, and whether their phase adds spatial information beyond the canonical population rate code. It is also unknown whether analogous phase tuning occurs for individual semantic concepts.

To test these questions, we first analysed human single-neuron recordings during virtual navigation and an openly available rat hippocampal linear-track dataset. In both datasets, we asked whether spikes showed location-specific theta-phase structure and repeated the analysis after excluding neurons with detectable rate place tuning. Finally, in the human navigation dataset, we asked whether phase features from neurons without a rate place label improved location decoding beyond firing rates from the full population. We then analysed human hippocampal neurons during a visual tuning task, asking whether individual images produced concept-specific phase structure and whether this remained after removing images that already showed firing-rate tuning for the same neuron.

In human and rodent navigation, location-specific phase tuning exceeded chance and remained after rate place cells were excluded. Most strikingly, in human navigation phase features from neurons without a rate place label improved location decoding beyond firing-rate features from the full population. In the visual task, concept-specific phase tuning also remained significant when rate-tuned images were removed from each neuron’s analysis.

Together, these findings show that theta-relative spike timing reveals spatial information beyond canonical rate labels and adds to the full population rate representation. The visual task extends this principle to concept-specific phase structure.

## Methods

We analysed single-neuron spiking and local field potential (LFP) phase in three datasets: the human CityBlock virtual-navigation dataset, the public HC3 rodent linear-track dataset (Mizuseki et al., 2013, 2014), and a human visual tuning dataset in which images were presented to epilepsy patients to identify putative concept neurons (Kolibius et al., 2023).

Across datasets, each retained spike was assigned an instantaneous theta phase from the spike-interpolated LFP. In CityBlock and HC3, spikes were assigned to spatial position bins and pooled across repeated loops through the virtual environment or traversals of the linear track, respectively. In the human visual tuning task (Kolibius et al., 2023), spikes were aligned to image onset and analysed within pre- and post-image windows. A separate population analysis tested whether post-image phase features decoded broad stimulus category.

A phase-tuned neuron was defined as a neuron that showed greater spike-phase concentration in at least one eligible spatial bin in CityBlock or HC3, or a greater phase-vector difference during image presentation versus baseline for at least one eligible image in the visual tuning task, than expected under its within-neuron null. We refer to these neurons as phase place cells in the spatial tasks and phase concept neurons in the semantic tuning task. In contrast, a rate place cell was defined as a neuron whose firing rate differed across spatial bins more strongly than expected under a rate-shift null.

Classical theta phase precession is typically observed in spatially rate-tuned neurons: as an animal traverses a place field, spikes occur at progressively earlier phases of the ongoing theta cycle. This creates a potential confound for analyses of phase-based spatial coding, because apparent phase tuning could partly reflect conventional rate place fields sampled at different theta phases. To test whether spatial phase tuning extended beyond neurons with detectable rate place tuning, we repeated the phase-tuning analyses after excluding rate place cells at increasingly liberal thresholds: no exclusion, p ≤ 0.05, and p ≤ 0.20. These exclusions were applied before phase testing, so the phase-testable denominator changed across exclusion thresholds. This effect was especially pronounced in HC3, where rate place tuning was common and rate place exclusion substantially reduced the tested denominator.

The single-neuron spatial analysis used pairwise phase consistency (PPC) as the primary phase-concentration statistic. PPC is a bias-corrected measure of phase consistency based on pairwise phase differences between spikes (Vinck et al., 2010). Under independent sampling from a fixed phase distribution, its expected value does not depend on spike count. We therefore preferred PPC over circular standard deviation when comparing bins with different spike counts. For each neuron, PPC was computed separately in each eligible spatial bin, and the observed neuron-level statistic was the maximum PPC across eligible bins. CityBlock population analyses used support-vector machine decoders with a radial basis function kernel (SVM-RBF) to compare spatial information carried by firing-rate features, phase features, and combined rate-plus-phase features. The concept neuron population decoder used post-image phase-vector features to classify image category.

## Datasets and recordings

### CityBlock

The analysed CityBlock subset comprised human intracranial single-neuron and LFP recordings from four recorded participants across six task sessions. This dataset comprised 192 microwire electrodes, which contained 236 putative single units.

Spike sorting was performed with Combinato (Niediek et al., 2016).

Participants navigated through a virtual environment (see Fig. 1) and were asked to remember the spatial positions of six items distributed through the environment on each loop. During item encoding, stimulation was delivered through a white-matter contact at 3 Hz or 8 Hz, or no stimulation was delivered (0 Hz/no stimulation). Because of technical errors, the present analysis was restricted to spatial position along the route.

**Figure 1.**
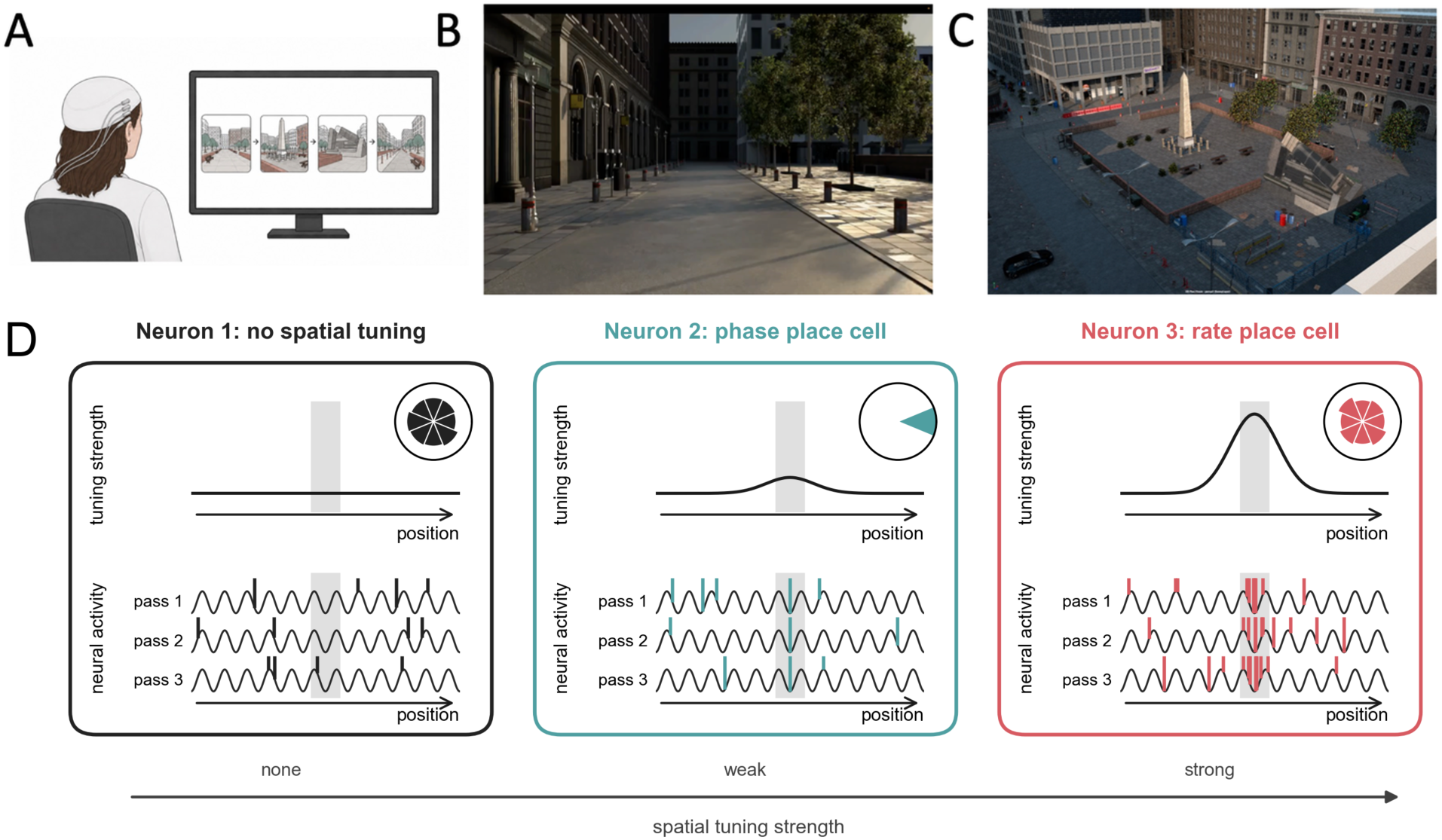
CityBlock task and schematic distinction between rate- and phase-based spatial tuning. **(A)** Participants navigated a virtual environment while hippocampal single-neuron activity and local field potentials were recorded with intracranial microwires. **(B)** Example first-person view from the CityBlock virtual environment. **(C)** Overhead view of the environment. (**D)** Schematic of three different neurons, ordered by spatial tuning strength. Top, tuning strength as a function of position along the route. Bottom, spikes (vertical lines) relative to the theta rhythm (3–12 Hz) during three passes along the whole route. The grey band marks the same location in each box, the tuned location of neurons 2 and 3. Neuron 1 (no spatial tuning) fires at baseline at random theta phases. Neuron 2 (phase place cell) shows spikes occurring close to its tuned location at a consistent theta phase. Neuron 3 (rate place cell) fires more when close to its tuned location. Circular histograms show the spike phases close to the tuned location. Schematics are conceptual.

Spiking activity was obtained from microwire recordings, and LFP phase was estimated from local microwire LFP signals after spike interpolation and artifact rejection. Loops were defined from the encoding trajectory intervals. The analysed sessions contained either 12 or 24 encoding loops.

CityBlock recording sessions acquired within 8 h of one another were concatenated for analysis because spike sorting was performed jointly across those sessions and the same isolated units were tracked across them. Intervening periods without active recording were treated as recording gaps. Accordingly, filtering and artifact-threshold estimation were performed separately within each contiguous recorded segment, so that discontinuities between segments did not influence thresholds in neighboring data.

### Rodent linear track (HC3)

The HC3 analyses used publicly available hippocampal recordings from rats running on linear tracks (Mizuseki et al., 2013, 2014). For the primary pooled four-bin phase-tuning analysis, 1,257 hippocampal units had sufficient spikes, distributed across sufficient traversals, to estimate phase tuning in at least one spatial bin. This set comprised 1,159 CA1 units, 91 CA3 units, and 7 dentate-gyrus units. According to the official HC3 cell-type annotations, 845 were putative pyramidal cells, 372 were putative interneurons, and 40 were unclassified. Because spatial-bin eligibility was recomputed after each circular-shift permutation, some neurons did not yield a valid null statistic on enough permutations for reliable inference. We therefore required each neuron to yield a valid null statistic in at least 75% of the 5,000 permutations. Nine hundred and fifty-six neurons met this criterion and entered the final pooled analysis: 885 from CA1, 64 from CA3, and 7 from dentate gyrus. Of these, 558 were annotated as putative pyramidal cells, 366 as putative interneurons, and 32 were unclassified.

LFP phase was estimated from the official HC3 local field potential recordings sampled at 1,250 Hz. For each unit, the local LFP channel was selected from the unit’s own electrode group by comparing spike waveforms across recording sites and choosing the site with the largest peak-to-peak median waveform amplitude.

### Human visual semantic tuning dataset

The visual semantic tuning dataset comprised hippocampal single-neuron recordings from 14 participants (7 female; age range, 19–58 years) across 20 recording sessions, yielding 216 putative single units. Participants were implanted with Behnke-Fried depth electrodes with microwire bundles, and hippocampal microwires were localized by MRI/CT co-registration. Data acquisition, spike detection, and spike sorting followed the procedures described previously (Kolibius et al., 2023).

During the visual tuning task, every image was presented six times in pseudorandom order. Each image was shown for 1 s with a 500–550 ms jittered inter-image interval, and participants categorized each image as an animal, place, or face to maintain attention during stimulus presentation.

Putative firing-rate concept neurons were identified from the visual tuning task by testing stimulus-evoked firing against pre-stimulus baseline activity. Because only pictures were presented, we could not test whether these responses were invariant across sensory modalities. We therefore refer to these neurons, and to the phase concept neurons described below, as putative concept neurons. For each unit and image, the 1 s post-image period was divided into 19 overlapping 100 ms bins. Spike counts in each bin across presentations of that image were compared with the 500 ms pre-image baseline periods pooled across all images in the session using two-tailed Mann-Whitney U-tests, with Simes correction for multiple comparisons. The conventional concept neuron threshold was P < 0.0005. These firing-rate concept neuron labels and image-level firing-rate tuning labels were used for overlap descriptions and sensitivity analyses.

Ethical approval was granted by the Ethik-Kommission of the Friedrich-Alexander Universität Erlangen-Nürnberg (142_12 B). Informed consent was obtained in accordance with the Declaration of Helsinki.

### LFP preprocessing, spike interpolation, artifact rejection, theta phase estimation, and low-theta-power exclusion

LFP preprocessing was designed to estimate theta phase while reducing direct contamination from action potentials. Before phase estimation, LFP samples around spike times were replaced by linear interpolation between the two LFP samples immediately outside the spike-centred replacement window. Overlapping replacement windows were merged before interpolation. This procedure generated the spike-interpolated LFP used for theta-phase estimation.

For CityBlock, preprocessing was performed on the native continuous microwire recordings sampled at 30,000 Hz. Spike-centred interpolation was applied at the native sampling rate using a 3 ms replacement window, after which the LFP was downsampled to 1 kHz. For HC3, spike interpolation was applied to the official 1,250 Hz LFP traces by replacing the three-sample window centred on the LFP sample corresponding to each spike. For the visual semantic tuning dataset, spike-centred interpolation was performed at the native sampling rate using the same 3 ms replacement-window rule as in CityBlock, followed by MNE-based resampling to 1 kHz.

Artifact-contaminated periods were identified before assigning theta phase values to spikes. Artifact detection was adapted from the robust multi-feature procedure described by Kunz et al. (2024) and applied to microwire LFPs by Guth et al. (2025), in which candidate artifacts are identified from complementary amplitude-, gradient-, and broadband-power-based features. Detection was performed on the spike-interpolated LFP within each contiguous recorded segment. The LFP was first bandpass filtered from 0.5 to 150 Hz using a fourth-order Butterworth filter. Candidate artifact samples were defined as the union of samples exceeding robust thresholds on any of the three features. Filtered-amplitude and temporal-gradient outliers were identified using two-sided median ± 4 × interquartile range thresholds, whereas broadband-power outliers were identified using a one-sided median + 4 × interquartile range threshold. Broadband power was summarized as the summed standardized log Morlet power across 30 log-spaced frequencies from 1 to 60 Hz, using seven-cycle wavelets. Candidate artifact intervals were padded by 250 ms on either side, merged when overlapping, and clipped to remain within the corresponding contiguous recorded segment. Samples within the resulting padded artifact mask were excluded from valid phase estimates and from occupancy or event-window denominators.

For each continuous recording segment, theta phase was estimated from the local spike-interpolated LFP using a theta-wave interpolation procedure adapted from Qasim et al. (2021), with a 3–12 Hz instead of a 2–10 Hz theta band. The LFP was first filtered in the 3-12 Hz theta band to define successive trough-peak-trough sequences. A broader 30 Hz low-pass filtered copy of the same LFP was then used to refine the timing of these landmarks and to identify the rising and falling voltage midpoints within each cycle. Troughs were assigned 0 and 360 degrees, rising midpoints 90 degrees, peaks 180 degrees, and falling midpoints 270 degrees. Phase was linearly interpolated between adjacent landmarks and then represented in radians on a circular scale. This yielded one theta-phase estimate for each LFP sample within complete theta cycles.

Theta phase was estimated from the spike-interpolated LFP for all three datasets: CityBlock, HC3, and the visual tuning dataset. Because phase estimates are less reliable when theta-band power is weak, CityBlock and the visual tuning dataset excluded samples in the lowest quartile of theta-band power within each contiguous real recording segment. HC3 did not apply the low-theta-power exclusion because theta oscillations dominate the rat hippocampal LFP during running (Buzsáki, 2002), and the HC3 analysis was restricted to running periods. Where applicable, artifact-masked and low-theta-power samples were excluded from both spike-phase assignment and occupancy or event-window denominators.

### Behavioral and event alignment

#### CityBlock spatial binning

For CityBlock, task coordinates were linearized within each encoding loop. Loop progress was defined as the cumulative Euclidean distance along the logged trajectory and was normalized to range from 0 to 1 within each loop. Samples or spikes occurring before the first logged coordinate or after the last logged coordinate were treated as unmapped and excluded from spatial feature construction. Valid mapped samples and spikes were assigned to one of 20 equisized spatial bins along the normalized coordinate.

Item-presentation and stimulation periods were excluded before assigning spike phases to spatial bins and before computing occupancy denominators and decoder features. Item-presentation periods were removed from item onset to item offset. Pre-item stimulation windows were removed relative to item onset depending on stimulation condition: 3 Hz trials used −3667 to −333 ms, and 8 Hz trials used −1375 to −125 ms. No pre-item stimulation window was removed for the 0 Hz/no stimulation condition.

#### HC3 linear-track binning

For HC3, official position samples were projected onto a one-dimensional track coordinate using the first principal component of the two-dimensional position trace and scaled to a 0–250 cm coordinate. The analysed core segment was restricted to 40–210 cm, excluding the first and last 40 cm of the track. These excluded track ends are reward regions where animals reduced their running speed (Parra-Barrero et al., 2021).

Speed was computed as absolute central-difference speed on the scaled one-dimensional coordinate, smoothed over 0.1 s, and interpolated to LFP samples. Samples were retained only when position was valid, the animal was within the analysed core segment, and speed was greater than 10 cm/s. The core segment was divided into 4 equal spatial bins. Complete passes through the core segment were classified by direction as increasing or decreasing along the scaled coordinate. Analyses were performed separately for pooled traversals, increasing traversals, and decreasing traversals.

### Visual semantic tuning image-event windows

For the visual tuning dataset, spikes were aligned to image onsets. Images were displayed for 1 s. Post-image phase structure was quantified from 0.3–1.0 s after image onset. The pre-image phase baseline for each neuron was the −0.5–0 s window before image onset, pooled across all images for the same neuron. Post-image and baseline spike counts were not downsampled to match. Only spikes occurring during valid, artifact-free, and non-low-theta-power LFP samples within these windows entered the spike-phase analysis.

### Feature construction and eligibility criteria

For every retained spike, the analysis stored the spike’s theta phase, the assigned behavioral bin, and the loop, traversal, or image-presentation identity. In CityBlock and HC3, a spike was retained for spatial analysis only if the corresponding LFP sample passed the real-sample, artifact, phase-validity, position-validity, and dataset-specific behavioral masks. In HC3, the primary mask was: real sample, not artifact, theta-wave-interpolated phase present, finite unmasked phase, valid position, within the 40-210 cm core interval, and speed greater than 10 cm/s. In the visual tuning dataset, a spike was retained for event-locked phase analyses only if it occurred within the specified pre-image or post-image window and its LFP sample passed the phase-validity, artifact, and low-theta-power masks.

Occupancy was defined as the amount of valid LFP time in a loop or traversal and spatial bin after all masks were applied. Firing rate was defined as spike count divided by this valid occupancy time. For CityBlock decoder analyses, rate features were loop-by-bin firing rates for each neuron. Phase features were the first circular moment of spike phase for each neuron and bin, represented by the mean cosine and mean sine of spike phases. The combined rate-plus-phase feature set concatenated the rate and phase features.

Before computing spatial phase tuning, each neuron was evaluated separately within each spatial bin. A neuron-position bin was included only when it had sufficient sampling across the relevant behavioral units: loops in the virtual environment for CityBlock and track traversals for HC3.

A bin was retained if three criteria were met. First, the neuron had to emit at least 10 spikes in that bin after all preprocessing and behavioral exclusions. Second, those spikes had to be distributed across at least half of the loops or traversals in which the animal or participant occupied that bin, rounded up to the nearest whole loop or traversal. Third, no single loop or traversal was allowed to contribute more than 50% of the spikes in that bin. This prevented phase-tuning estimates from being driven by a single unusually active pass through a location.

The primary CityBlock analyses divided the environment into 20 spatial bins. The primary HC3 analyses divided the retained 40–210 cm track segment into 4 equal spatial bins. The same bin-level eligibility criteria were used for the single-neuron PPC analyses and population decoding. For population decoding, a neuron was included only if at least three spatial bins met the spike-count, loop/traversal-coverage, and single-pass dominance criteria described above. Thus, decoder-eligible neurons had at least 3 of 20 eligible bins in CityBlock and at least 3 of 4 eligible bins in HC3.

For the visual semantic tuning phase analysis, eligibility was defined at the neuron-image level. An image was eligible for a neuron if the post-image analysis window contained at least 10 valid spike phases, those spikes occurred on at least 3 of the 6 image presentations, no single presentation contributed more than 50% of the post-image valid spikes for that image, and the post-versus-baseline phase statistic was finite. A neuron was phase-testable if it had at least one eligible image.

### Single-neuron spatial phase tuning

Single-neuron phase tuning was quantified with PPC. For a given neuron and bin, PPC was computed from the phases of retained spikes assigned to that bin. The observed phase-tuning statistic for a neuron was the maximum PPC across all eligible bins in the relevant configuration and direction mode. If a neuron had no eligible bins, it was not phase-testable for that configuration.

Statistical significance was assessed with a phase-shift null. For each permutation, spatial-bin labels were circularly shifted within each loop or traversal by a nonzero offset. The offsets were sampled independently across loops or traversals. After shifting, bin eligibility and PPC values were recomputed, and the maximum PPC across eligible bins was stored as one null sample. The per-neuron phase-tuning p-value was computed as:

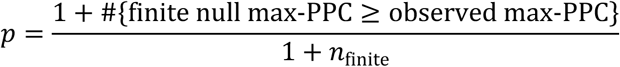

Here, *n*_finite_ is the number of finite null max-PPC values for that neuron. Neurons were classified as phase-tuned at neuron-level alpha 0.05. CityBlock and HC3 single-neuron analyses used 5,000 permutations and required at least 75% of phase-shift permutations to yield a finite max-PPC statistic after spatial-bin eligibility was recomputed. Sparse null draws were not resampled.

Across 3,876 HC3 source records, 956 were phase-testable in pooled traversals, 1,012 in increasing traversals, and 973 in decreasing traversals before rate place exclusion. Most exclusions resulted from insufficient eligible spatial bins, with additional exclusions due to insufficient finite-null support.

### Rate place testing and rate place exclusion

Rate place tuning was assessed separately from phase tuning. For each neuron, loop/traversal firing rates were computed from spike counts and valid occupancy. The rate place statistic was a one-way ANOVA F statistic over spatial bins. A rate place null distribution was generated by independently circularly shifting each loop/traversal rate vector across spatial bins; unlike the phase-shift null, zero-bin shifts were allowed. The rate place p-value used the same add-one permutation rule as the phase-tuning p-value.

Rate place exclusion was used to define analysis populations in which phase tuning was evaluated after removing neurons that met a firing-rate spatial tuning criterion. A rate place cell was defined for a given dataset, configuration, and direction mode as a neuron with a rate place p-value at or below the specified threshold. Exclusion thresholds were either no exclusion, p ≤ 0.05, and p ≤ 0.20. Because rate place status was computed within each configuration and direction mode, rate place exclusion changed the denominator for each corresponding phase-tuning analysis.

For the additive population decoder, the rate place threshold of p ≤ 0.05 controlled which neurons contributed phase features. The rate-only feature matrix retained all decoder-eligible rate features, whereas the rate-plus-phase matrix added phase features only from neurons not excluded by the rate place threshold of p ≤ 0.05.

### Group-level inference for single-neuron phase tuning

Group-level inference tested whether the observed number of phase-tuned neurons exceeded the number expected under the phase-shift null. For every phase-testable neuron, the finite null max-PPC samples were converted into null p-values using the same add-one procedure used for the observed statistic. For each permutation index, the analysis counted how many neurons had null p-values at or below the neuron-level alpha. This produced a null distribution of tuned-neuron counts that preserved the per-neuron null structure and the same eligibility rules used for the observed analysis. In CityBlock and HC3, invalid per-neuron null draws were treated as missing for that neuron and permutation index after the 75% finite-null support rule was applied and contributed no tuned neuron to that permutation-index count; the group draw itself was retained. The group test used all 5,000 null counts, whereas each neuron-level p-value used its own finite-null denominator. The group p-value was the add-one one-sided probability of a null count greater than or equal to the observed tuned-neuron count.

### Population decoding

CityBlock population decoding tested whether phase features contributed to spatial-bin classification under the same behavioral and feature-validity rules used for the single-neuron analyses. Decoder analyses used an SVM-RBF model. Feature sets were rate, phase, and rate plus phase. Rate features were firing rates for each neuron and bin. Phase features were mean cosine and mean sine of spike phase for each neuron and bin. The combined feature set concatenated the rate and phase columns. Additive phase-shift decoder analyses contrasted the rate-only feature set with the rate-plus-phase feature set.

Cross-validation was grouped by loop, so that rows from the same loop were not split across train and test folds. CityBlock decoder analyses required at least 3 eligible spatial bins per included unit. For each session, analysis configuration, and rate place exclusion condition, 10 outer train/test splits were drawn over loops with a 0.75 training-group fraction. Accepted splits contained all spatial classes in both train and test sets. RBF hyperparameters were selected within the training data using 3 grouped inner folds over loops and the grid C = [0.1, 1.0, 10.0], γ = [“scale”, 0.01, 0.1, 1.0]. Training rows were downsampled to equalize class counts before fitting the classifier. Features were standardized using training-fold statistics only, and the same transformation was applied to the corresponding test fold. Nonfinite standardized feature values were set to zero after training-fold standardization.

Decoder inference used normalized macro-averaged mean squared error (MSE), with equal weight for each true spatial bin. Wrapped distances were divided by half the number of bins before squaring. For descriptive effect sizes, we also reported macro-averaged absolute wrapped error in spatial-bin units. CityBlock decoder errors used the wrapped distance on the linearized path.

Standalone spatial decodability of each feature set was assessed with a circular label-shift null. For each permutation, spatial-bin labels were circularly shifted within each loop while the feature matrix was kept fixed. The phase contribution was tested with a phase-shift decoder null. For each *phase-shift* permutation, phase-feature columns were circularly shifted across spatial bins within each loop by a nonzero offset, while rate features and spatial-bin labels were kept fixed. The decoder was then refit on the shifted feature matrix. Additive phase gain was defined as:

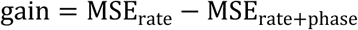

For each session, gains were averaged across the 10 outer splits; group inference averaged session gains. Positive values of this statistic indicate lower error for the rate-plus-phase decoder than for the rate-only decoder. Decoder p-values were computed with add-one permutation tests using the appropriate one-sided tail for the predefined statistic. CityBlock decoder analyses used n_perm = 5,000. Only SVM-RBF decoders with downsampled training classes were used for the CityBlock decoder analyses. The concept neuron category decoder is described separately below because it used image rows and category labels rather than spatial-bin rows.

The relative descriptive gain was the observed reduction in wrapped-bin error divided by the rate-only decoder’s improvement over its own label-null mean:

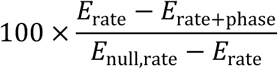

The rate-only label-null mean was the reference for this effect-size display; inferential significance was assessed separately with the phase-shift null. Shifted phase features can worsen decoder performance, so that null gain need not be centered on zero.

A separate phase-only decoder used neurons that did not meet the rate place criterion and was tested against the circular label-shift null.

### Visual semantic category decoding

The visual semantic population decoder tested whether post-image spike phase features decoded the broad stimulus category of each image. Categories were animal, face, and place, as defined by the stimulus identifiers in the original task design. The decoder was run separately within each recording session. Each row of the decoder matrix was one unique image in that session, collapsed across the six presentations of that image.

For each neuron and image, post-image spikes from 0.3 to 1.0 s after image onset were retained if their LFP samples passed the final phase-validity mask. A neuron-image pair contributed phase features if it met the same primary image-level eligibility criteria used for the single-neuron concept neuron phase analysis: at least 10 valid post-image spikes, spikes on at least 3 presentations, no single presentation contributing more than 50% of the post-image spikes, and finite PPC. A neuron contributed to the session decoder if it had at least 3 eligible images in that session.

The decoder used phase features only. For every eligible neuron-image pair, the feature vector contained the mean cosine and mean sine of the post-image spike phases. Ineligible neuron-image pairs were retained as missing values until model fitting. The decoder was run in two matched branches: one benchmark rate-image-excluded branch (the primary category-decoding analysis) and one sensitivity branch with no benchmark firing-rate image exclusion. In the rate-image-excluded branch, benchmark firing-rate tuned images were excluded from the phase feature for the same neuron. This exclusion operated at the neuron-image level: the corresponding feature was treated as missing for that neuron and image, but the neuron was not removed from the session, and the image row was not removed from the decoder.

Train/test splits were image-level and stratified by category. For each decodable session and repeat, 75% of image rows were assigned to training and 25% to testing, with train and test sets required to contain all three categories. Ten independent split repeats were run. Within each training split, rows were downsampled to equalize category counts before fitting the classifier. Features were standardized using training rows only. Column means and standard deviations were computed from finite training values. Nonfinite standardized values in the train and test matrices were then set to zero.

The classifier was a support-vector machine with a radial basis function kernel. Hyperparameters were selected by 3-fold stratified cross-validation within the training set using C = 0.1, 1.0, or 10.0 and gamma = scale, 0.01, 0.1, or 1.0. The performance metric was balanced accuracy, averaged across categories. Statistical significance was assessed with a category-label null. For each session and permutation, category labels were shuffled across image rows while preserving the number of images in each category. The feature matrix, missingness pattern, train/test split procedure, class downsampling rule, hyperparameter search, and scoring metric were otherwise unchanged. The decoder used 5,000 label permutations per session. For each session, observed balanced accuracy was averaged across the 10 real split repeats; each session-level null value was the corresponding mean balanced accuracy under a shuffled-label run.

Group-level decoder inference averaged observed balanced accuracy across decodable sessions. To form the group null, one shuffled-label score was sampled from each decodable session’s null distribution and the sampled scores were averaged across sessions. This was repeated 5,000 times. The group p-value was

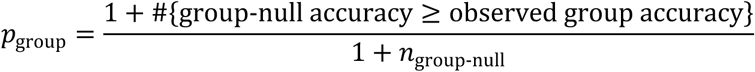

Sessions were excluded if they contained no eligible neurons, lacked sufficient category representation, or could not support the required inner cross-validation. Excluded sessions were not assigned chance-level performance. In the present dataset, 18 of 20 sessions were decodable; two excluded sessions contained no eligible neurons.

### HC3 region and cell-type analyses

We compared phase place fractions across HC3 regions and cell types using the official HC3 annotations. Both comparisons used pooled traversals and the phase-testable population before rate place exclusion. The regional comparison was restricted to CA1 and CA3. Dentate gyrus was omitted because only seven phase-testable neurons were available. The cell-type comparison retained official pyramidal/interneuron annotations and excluded 32 unclassified units. Each comparison used a two-sided Fisher exact test of the counts of phase place cells and other neurons. These exploratory association tests were performed at the neuron level, with each phase-testable neuron contributing one observation; without modelling clustering within sessions or animals.

### Rate/phase overlap analyses

We summarized rate place fractions separately among phase place cells and neurons without detected phase place tuning in CityBlock and in pooled HC3 traversals. Rate place status was defined at p ≤ 0.05, and phase place status by the PPC phase-shift test at neuron-level alpha 0.05. Both groups were restricted to phase-testable neurons with finite rate place p-values under the current 5,000-permutation, 75% finite-null-support analysis. Phase-ineligible neurons were excluded from both denominators. These are descriptive classification-overlap comparisons using the same spike trains and behavioral masks as the primary analyses.

For the visual tuning dataset, benchmark firing-rate concept neuron annotations and benchmark firing-rate tuned images were used for overlap and sensitivity analyses. The inclusive phase concept analysis retained all candidate images. The rate-image-excluded sensitivity removed benchmark firing-rate tuned images from the phase concept candidate set at the neuron-image level. This sensitivity tests whether phase concept evidence persists after excluding images already labelled as firing-rate tuned for the same neuron. It does not test a denominator in which all benchmark firing-rate concept neuron units are removed.

### Visual semantic tuning event-locked spike-phase analysis

For the visual tuning dataset, spike phases were analysed in event-locked windows around image presentation. After LFP preprocessing, spike interpolation, artifact rejection, theta-phase estimation, and low-theta-power exclusion, spikes were assigned to either a post-image window or a pre-image baseline window. The post-image phase window spanned 0.3-1.0 s after image onset. The baseline was defined as the −0.5 to 0 s period before image onset, pooled across all images for the same neuron. Thus, the retained event-locked analysis compared post-image phase structure with a collapsed pre-image baseline from the same neuron. It did not match post-image and baseline window durations and did not downsample spikes to match pre/post spike counts.

For each neuron and image, we quantified the difference between the post-image and pre-image spike-phase distributions using a bias-corrected squared distance between their circular phase vectors. For a set of *n_A_* spike phases *θ_j_*, we first defined the summed phase vector as:

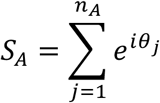

Phase consistency within set *A* was quantified using pairwise phase consistency (PPC):

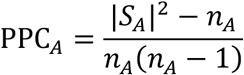

For two phase sets *A* and *B*, their cross-pair phase consistency was:

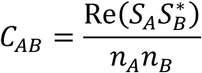

where 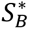 denotes the complex conjugate of *S_B_*. The post-versus-baseline distance was then:

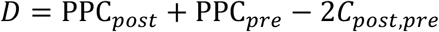

Larger values of *D* indicate greater separation between the post-image and pre-image phase distributions.

An image was included only if the post-image analysis window contained at least 10 valid spike phases, spikes occurred on at least 3 out of 6 image presentations and no single presentation contributed more than 50% of the post-image valid spikes for that image. Significance was assessed with a spike-label shuffle in which the combined post-image and collapsed pre-image phase pool was randomly relabelled while preserving the observed post-image and baseline spike counts. We used 5,000 permutations. For each image, the permutation p-value was calculated as

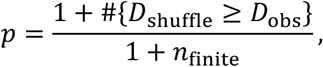

where *D*_obs_ is the observed post-versus-baseline distance and *n*_finite_ is the number of permutations that yielded a finite shuffled statistic. An image was classified as phase significant at p <= 0.001. A neuron was classified as a phase concept neuron if it had at least one eligible image with p <= 0.001. For group-level inference, each permutation generated a null count of phase concept neurons using the same eligibility and image-level threshold. The group p-value was the add-one probability of observing at least as many phase concept neurons under this null count distribution as in the real data. The primary concept neuron single-neuron rerun required at least 75% of the 5,000 permutations to yield finite statistics for every observed-eligible image in the tested neuron. If this finite-null requirement was not met, that neuron was excluded from the testable denominator rather than being retained with insufficient permutation evidence.

In the main analysis, images that already showed firing-rate tuning for a given neuron were excluded from that neuron’s phase concept analysis. This exclusion was applied at the neuron-image level: the neuron remained in the analysis, and the same image remained eligible for other neurons if it was not rate-tuned for them. The collapsed pre-image baseline pool was unchanged. We also ran an inclusive analysis in which these rate-tuned images were retained.

### CityBlock preferred phase and within-neuron sampling comparisons

Within phase place cells, we compared preferred phases in phase-tuned spatial bins with those in the same neurons’ eligible untuned bins. Tuned bins were those whose PPC exceeded the neuron’s max-statistic null at the 0.05 threshold. Preferred phase was the angle of the summed spike-phase vectors within each bin. The test statistic was the absolute wrapped difference between the circular means of the tuned and untuned bin phases, with each bin contributing one unit vector. We generated 100,000 permutations by shuffling bin labels within neuron while preserving each neuron’s tuned-bin count. Confidence intervals used 10,000 bootstrap resamples of neurons, retaining each neuron’s complete set of bins. We report the signed tuned-minus-untuned difference as the effect size. Analyses were repeated after excluding rate place cells at p ≤ 0.05.

We also compared valid occupancy time and firing rate between these bin sets. Firing rate was total retained spikes divided by valid occupancy in each bin. Each metric was standardized across eligible bins within neuron, and each neuron’s tuned-minus-untuned mean difference was then averaged with equal weight across neurons. Two-sided p-values used 100,000 within-neuron label permutations preserving tuned-bin counts. Confidence intervals used 10,000 neuron bootstrap resamples. These post hoc comparisons were not adjusted for multiplicity. Preferred-phase sensitivity simulations used 1,000 datasets per imposed circular shift and 1,000 permutations per simulated dataset.

### CityBlock phase precession

We tested whether spike phase changed systematically with position around each neuron’s preferred location, following Qasim et al. (2021). The analysis included all CityBlock phase place cells and rate place cells, including rate place cells that were not phase-testable, and used the same spikes, theta phases, and masks as the single-neuron phase analysis. The reference location was the centre of the maximum-PPC bin for phase place cells and the centre of the bin with the highest firing rate for rate place cells. Firing rate was computed as total spikes divided by total valid occupancy across loops. Neurons that met both criteria were analysed in both groups. We retained spikes within 1.5 bins of the reference location, that is, in the reference bin and in one neighbouring bin on each side. Each spike’s position was its signed distance from the reference location along the direction of travel. As in Qasim et al. (2021), each loop contributed at most the 11 retained spikes closest to the reference location. Neurons with fewer than 10 retained spikes were not tested. We fitted the circular-linear regression of Kempter et al. (2012) to spike phase and position, with the slope restricted to at most 1.5 theta cycles across the range of retained positions, and used the circular-linear correlation, signed by the fitted slope, as the test statistic. The null distribution was generated by redrawing each spike’s phase with replacement from all of the neuron’s analysed spike phases while keeping positions fixed (5,000 permutations). A neuron showed significant phase precession when its correlation fell in the lower tail of this null distribution (one-sided add-one permutation p ≤ 0.05), that is, when its spikes occurred at progressively earlier theta phases as the participant moved through the reference location. Phase changes in the opposite direction were tested with the upper tail. For each group, a one-sided binomial test compared the number of neurons with significant phase precession with the 5% expected by chance. A two-sided Fisher exact test compared the proportion of precessing neurons between phase place cells and rate place cells that were not phase place cells.

### HC3 arrival-end enrichment

We tested post hoc whether the maximum-PPC bin of phase place cells occurred disproportionately at the arrival end of a traversal. Within the four-bin 40–210 cm track segment, arrival was bin 4 for increasing traversals and bin 1 for decreasing traversals. For each significant neuron and direction, the null assigned its best bin uniformly among that row’s eligible bins. We generated 100,000 independent randomizations, preserving the number of significant rows and their eligibility masks, and used add-one one-sided p-values for arrival-bin counts. The two directions were evaluated with and without exclusion of rate place cells at p ≤ 0.05. To test whether this enrichment followed the location of place fields, we applied the same test to rate place cells, defined separately for each traversal direction (rate place p ≤ 0.05). For each rate place cell and direction, the tested bin was the bin with the highest firing rate, computed as total spikes divided by total valid occupancy across that direction’s traversals, and the null placed this bin uniformly among the bins with valid occupancy. These post hoc tests were unadjusted and did not model clustering within sessions or animals. Locations are described as arrival ends of the analysed segment, without assuming reward-zone identity.

### Statistical conventions and reproducibility

Permutation tests used fixed random seeds and add-one p-values. Unless stated otherwise for a specific statistic, one-sided permutation p-values were computed in the direction of the predefined observed statistic. Eligibility, rate place exclusion, phase-tuning statistics, and decoder feature construction were recomputed inside permutation procedures whenever the permutation changed the spike-to-bin, spike-to-window, feature-to-label, or feature-to-spatial-bin assignment.

## Results

Across datasets, we tested whether theta phase carries task information that is not fully captured by firing rate. We first tested spatial phase tuning in the CityBlock human virtual-navigation task (Fig. 1). We then applied the same spatial framework to hippocampal recordings from rats running on a linear track (Fig. 4). Finally, we tested whether neurons showed phase tuning to individual concepts in a visual concept tuning task (Fig. 5). Together, we use these analyses to test whether spike timing within the theta cycle carries structured information across human navigation, rodent navigation, and semantic representation.

### Theta phase tracks position in human virtual navigation

CityBlock was the human spatial dataset. We tested whether individual neurons fired at consistent phases of the theta rhythm at particular positions in the virtual environment (example neuron in Figure 2). A spatial bin entered the phase analysis only when the neuron fired enough spikes there, when spiking was observed on at least half of valid route loops, and when no single loop contributed more than half of the spikes. These constraints make the test less vulnerable to isolated bursts or inconsistent firing in few laps. Because phase precession in a rate-defined place field can appear as phase locking in neighboring spatial bins, we also repeated the analysis after removing neurons with firing-rate place fields.

**Figure 2.**
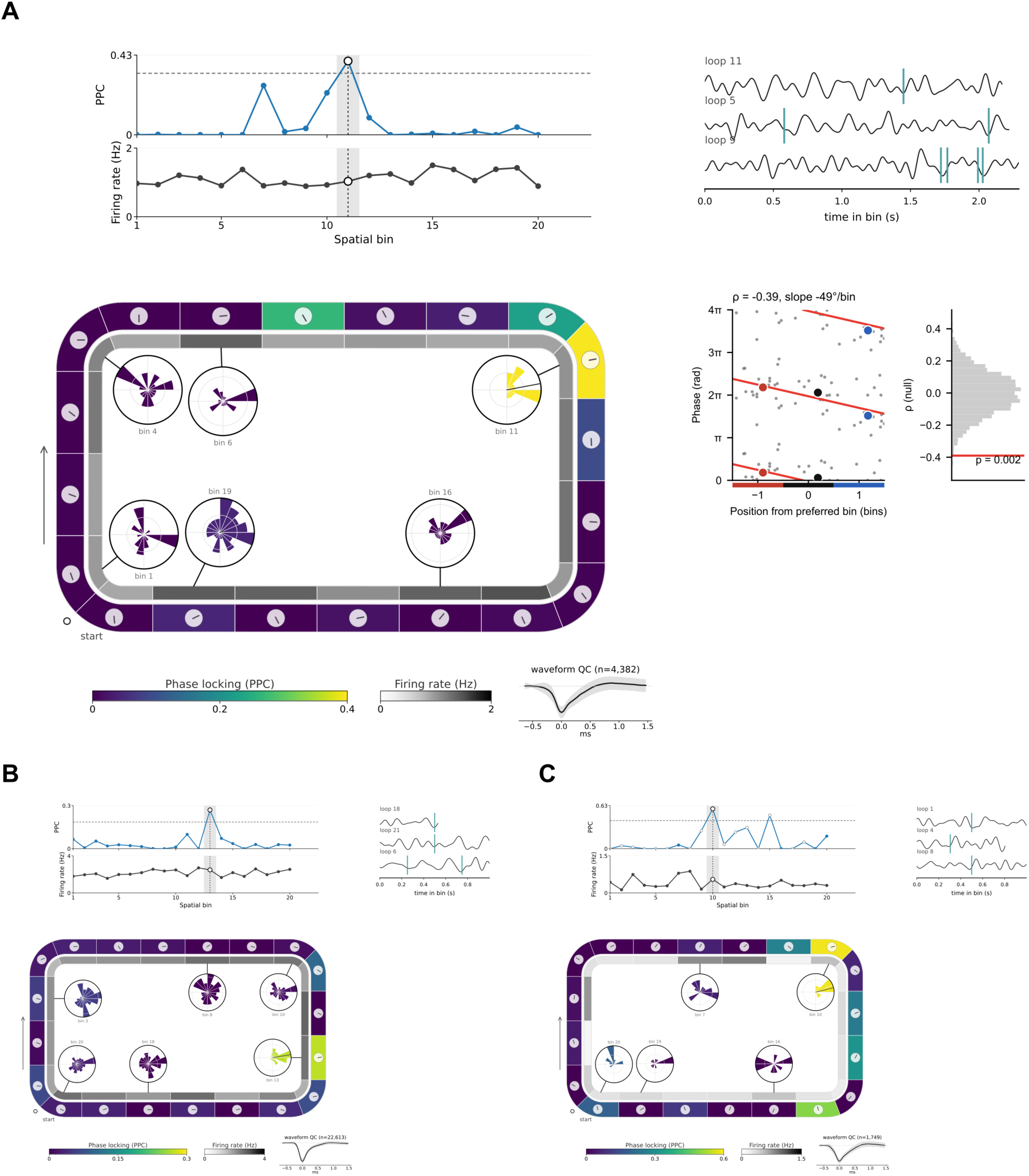
Example location-specific phase tuning in CityBlock neurons without detectable rate place tuning. **(A–C)** Three example hippocampal neurons. Top left, linearized pairwise phase consistency (PPC, blue) and firing rate (black) across the 20 spatial bins of the route. The dashed line marks the max-statistic permutation threshold (p = 0.05), and the gray band and open black circle mark the bin with the strongest phase locking. Open gray circles mark bins that were not tested because they contained too few spikes. Bottom, the route colored by PPC (outer band) and by firing rate (inner band). Small dials along the route show the preferred theta phase in each bin, and circular histograms show spike-phase distributions for selected bins. Top right, three passes through the bin with the strongest phase locking, showing the 3–12 Hz LFP and the neuron’s analysed spikes (teal), over the whole pass in A and over 1 s around the spikes in B and C. Waveforms show the mean spike waveform and its variability. Arrows indicate the direction of travel. (A, middle right) The neuron in A also showed phase precession. Theta phase of each spike is plotted against its position relative to the preferred bin over two cycles, with the circular-linear fit (red) and the mean phase of early, midpoint and late spikes (colored dots). The adjacent histogram shows the null distribution of the circular-linear correlation, with the observed value in red (Supplemental Figure 2).

CityBlock contained more neurons phase tuned to specific locations than expected under the shuffled null. Without excluding rate place cells, 14 of 153 phase-testable neurons showed above-chance phase locking to at least one spatial bin (null mean 7.5 neurons; *p*=0.0196). After excluding rate place cells, the effect remained: 12 of 132 phase-testable neurons were phase place cells (null mean 6.5; *p*=0.0322) (Fig. 2). Thus, a subset of CityBlock neurons carried location-specific theta phase information even after removing neurons with firing-rate spatial tuning (Figure 3).

**Figure 3.**
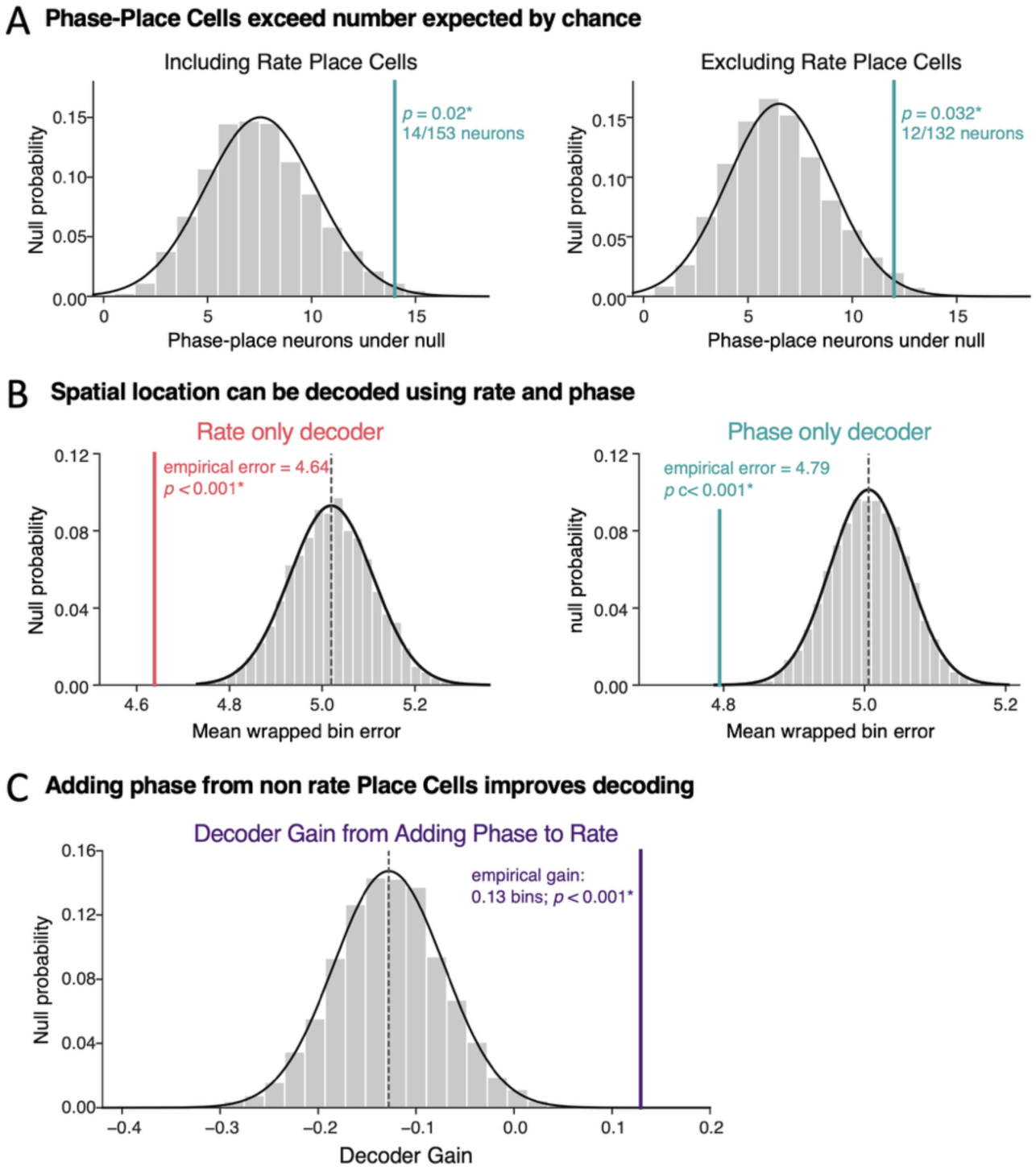
Theta phase carries spatial information beyond rate place coding in CityBlock. **(A)** The number of phase place cells exceeded that expected under the phase-shift null both when all phase-testable neurons were included (14/153 neurons; *p* = 0.020) and after neurons with significant rate place tuning were excluded (12/132 neurons; *p* = 0.032). Grey histograms show null distributions of phase place cell counts; teal lines indicate the observed counts. **(B)** Spatial position could be decoded independently from firing-rate and theta-phase features. The rate-only decoder achieved a mean wrapped error of 4.64 spatial bins, and the phase-only decoder an error of 4.79 bins; both were lower than their respective circular label-shift null distributions (*p* < 0.001). Lower error indicates better decoding. **(C)** To test whether phase contributed spatial information beyond the population firing-rate code, firing-rate features from the full population were held fixed while phase features were added only from neurons without detectable rate place tuning. Adding these phase features reduced mean wrapped decoding error by 0.13 spatial bins. This improvement exceeded the paired phase-shift null (*p* < 0.001), showing that theta phase in neurons without detectable rate place tuning carried spatial information not captured by population firing rates alone. In **(B–C)**, grey histograms show permutation-null distributions, dashed lines indicate their means, and colored vertical lines indicate the observed values.

Among phase-testable CityBlock neurons, 2 of 14 phase place cells (14.3%) met the rate place criterion, compared with 19 of 139 neurons without detected phase place tuning (13.7%). Phase-ineligible neurons were excluded from both denominators. In none of the three datasets did neurons with phase tuning but no rate tuning fire significantly less than neurons with rate tuning but no phase tuning (Supplementary Results).

Firing rate and theta phase each carried spatial information at the population level. The rate-only decoder performed above its shuffled control (mean wrapped error 4.64 vs. 5.02 bins; *p* = 0.00019996), as did the phase-only decoder (4.79 vs. 5.01 bins; *p* = 0.00039992). Phase-only and rate-only decoding errors did not differ significantly (paired sign-flip test across six sessions, p = 0.13). Phase features could improve the rate decoder without outperforming firing rates, provided they carried spatial information that the firing rates lacked. We next asked whether phase carried spatial information that was not already available in the population firing-rate code. We therefore kept firing-rate features from the full population fixed and added phase features only from neurons that did not meet the rate place criterion. Thus, any improvement had to come from spike timing in neurons that conventional rate place analysis would leave unlabeled. Across six sessions, the reduction in decoding error contributed by these phase features was 34.0% as large as the entire above-null advantage of the rate-only decoder. In absolute terms, mean wrapped error decreased by 0.13 spatial bins, from 4.64 to 4.51 bins. This improvement exceeded the paired phase-shift null (*p* = 0.00019996). Thus, theta phase from neurons without detectable rate place tuning carried spatial information that was not captured by firing rates from the full population (Figure 3). The information carried by phase and rate was therefore at least partly nonredundant.

### CityBlock preferred phase and sampling comparisons

Preferred phase did not differ detectably between phase-tuned and eligible untuned bins within phase place cells (24.09° versus 14.97°; signed difference 9.13°, 95% neuron-cluster-bootstrap CI −13.43° to 29.55°; within-neuron permutation p=0.764; 14 neurons, 15 tuned and 212 untuned bins). After excluding rate place cells, the difference was 5.28° (28.97° versus 23.68°; CI −21.73° to 30.60°; p=0.868; 12 neurons, 13 tuned and 174 untuned bins). These results do not establish equivalent preferred phases. Simulations indicated limited sensitivity to modest shifts: without rate place exclusion, estimated power was 16.7% at 20° and 28.9% at 30°, rising to 77.5% at 60°.

Within these neurons, firing rate did not differ detectably between phase-tuned and eligible untuned bins (mean standardized difference = −0.212, 95% neuron-bootstrap CI −0.814 to 0.361; p = 0.508; after rate place exclusion, −0.139, CI −0.831 to 0.514; p = 0.693). Occupancy was lower in phase-tuned bins (−0.586, CI −1.052 to −0.032; p = 0.061; after exclusion, −0.724, CI −1.144 to −0.233; p = 0.035). Confidence intervals and p-values derive from neuron bootstrap and within-neuron permutation procedures, respectively.

### Phase precession around the preferred location

In rate place cells, spike phase changes systematically with position through phase precession (O’Keefe and Recce, 1993; Qasim et al., 2021). We therefore asked whether spike phase also changed with position around the preferred location of phase place cells, using the approach Qasim et al. (2021) applied to phase precession in rate place cells (Methods). Four of 14 phase place cells showed significant phase precession (4/14 = 0.29), more than the 5% expected by chance (one-sided binomial test, p = 0.004, Supplemental Figure 2). All four were phase place cells without rate tuning, and none of the 14 showed a significant phase change in the opposite direction. Two of 24 rate place cells showed significant phase precession (2/24 = 0.083), which did not exceed chance (p = 0.34). At the prevalence that Qasim et al. (2021) reported (35/296 = 0.12), about three precessing rate place cells would be expected. The proportion of precessing neurons did not differ significantly between phase place cells and rate place cells that were not phase place cells (4/14 vs 2/22, two-sided Fisher exact test, p = 0.18). Thus, a significant number of phase place cells without rate tuning showed precession-like phase changes around their preferred location.

### Spatial phase tuning in rodent linear track

HC3 provided a rodent spatial benchmark for the same single-neuron phase place test. We restricted the analysis to the central portion of the track to avoid any confounds close to the reward sites, and to running periods, then tested whether hippocampal neurons fired at consistent theta phases at particular positions. As in CityBlock, bins had to contain enough spikes distributed across traversals rather than being dominated by one pass. This makes the comparison useful as a spatial phase-tuning benchmark (Figure 4).

**Figure 4.**
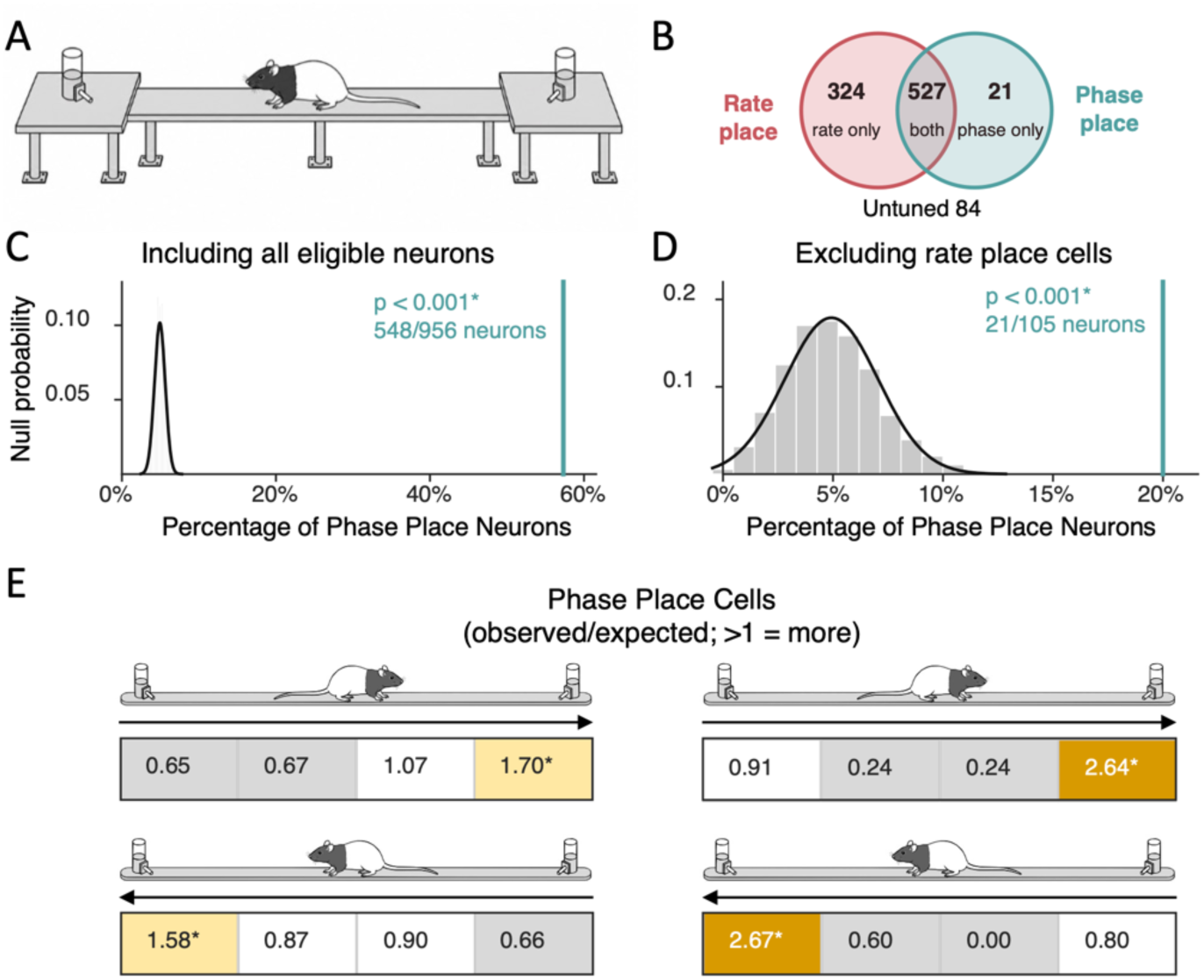
Spatial phase tuning in the rodent linear-track hippocampal dataset. **(A)** Rats traversed a linear track between the two ends of the environment. **(B)** Overlap between rate place and phase place classifications among the 956 phase-testable neurons in pooled traversals: 324 neurons were rate place only, 527 showed both rate- and phase place tuning, 21 were phase place only, and 84 showed neither form of tuning. **(C)** Phase place cells were substantially more frequent than expected under the phase-shift null when all phase-testable neurons were included (548/956 neurons; p < 0.001). **(D)** Phase place cells also exceeded the null expectation after rate place cells were excluded (21/105 neurons; p < 0.001), showing that location-specific phase tuning was not restricted to neurons with detectable rate place fields. In (C–D), grey distributions show the permutation null and teal lines indicate the observed fraction of phase place cells. (E) Phase place cells were not distributed uniformly along the analysed track segment. For each phase place cell and traversal direction, the spatial bin with the highest PPC was identified and its observed frequency was compared with that expected from the neuron’s eligible bins. Values show observed/expected ratios for each of the four spatial bins; values greater than 1 indicate enrichment. The top and bottom rows show the two traversal directions, indicated by arrows. The left column includes all phase place cells and the right column excludes rate place cells. In both directions, the strongest phase locking occurred disproportionately in the final spatial bin reached by the animal, both before rate place exclusion (1.70 and 1.58) and after rate place cells were excluded (2.64 and 2.67). Asterisks indicate significant enrichment.

HC3 showed widespread spatial phase tuning. In pooled traversals, 548 of 956 eligible neurons showed phase locking to at least one spatial bin (*p* = 0.00019996). The effect also appeared when traversals were split by running direction: 508 of 1012 neurons in increasing traversals and 471 of 973 neurons in decreasing traversals were phase place cells (both *p* = 0.00019996). Thus, the phase place analysis recovered a strong spatial phase signal in rodent hippocampus (Figure 4).

Among phase-testable HC3 neurons in pooled traversals, 527 of 548 phase place cells (96.2%) also met the rate place criterion, compared with 324 of 408 neurons without detected phase place tuning (79.4%). This overlap shows that rate place labels were common among phase place cells, while the residual analysis above identifies phase tuning among neurons without those labels.

The critical control was whether this signal survived after removing neurons with spatial firing-rate fields. That filter leaves a much smaller population, so the result should be read as evidence for a residual phase signal among neurons without detected rate place tuning. After excluding rate place cells, 21 of 105 eligible pooled-traversal neurons were phase place cells (*p* = 0.00019996); the direction-specific analyses also remained above chance (18 of 102 increasing, *p* = 0.00039992; 19 of 97 decreasing, *p* = 0.00019996). Thus, HC3 supports spatial phase tuning beyond rate place labels (Figure 4).

Because the eligibility criteria were stringent, only 956 of 3,876 HC3 source records were phase-testable in pooled traversals before rate place exclusion. The residual phase place result therefore reflects an excess among analyzable neurons that are not rate place cells, not the prevalence of phase tuning across all recorded units.

### Enrichment of phase place cells near reward zones

We next asked whether the location showing the strongest phase locking was systematically related to traversal direction. For each phase place cell, we identified the eligible spatial bin with the highest PPC and tested whether this bin occurred disproportionately at the arrival end of the analysed track segment, the final spatial bin reached in each traversal direction. In increasing traversals, 194 of 508 phase place cells had their maximum-PPC bin at the arrival end, compared with 114.17 expected under the null (observed/expected = 1.70, p = 0.000010). In decreasing traversals, 178 of 471 cells showed the same pattern (expected = 112.58, observed/expected = 1.58, p = 0.000010). The enrichment became stronger after excluding rate place cells. In increasing traversals, 11 of 18 phase place cells had their maximum-PPC bin at the arrival end (expected = 4.17, observed/expected = 2.64, p = 0.000350), and in decreasing traversals 12 of 19 did (expected = 4.50, observed/expected = 2.67, p = 0.000240). This enrichment could reflect place fields that cluster at the arrival end. We therefore applied the same test to rate place cells, using the bin with the highest firing rate in each traversal direction. In increasing traversals, 354 of 1,624 rate place cells had their peak firing-rate bin at the arrival end, compared with 406.00 expected under the null (observed/expected = 0.87, p = 0.999). In decreasing traversals, 428 of 1,597 rate place cells did (expected = 399.25, observed/expected = 1.07, p = 0.053). Thus, rate place cells did not show the arrival-end enrichment of phase place tuning, apart from a weak trend in decreasing traversals.

We next asked whether phase place tuning differed across hippocampal cell types and subregions. We used the pooled phase-testable HC3 population before rate place exclusion and excluded 32 units without a pyramidal or interneuron classification. Phase place cells were more often pyramidal cells than neurons without phase place tuning (365/543 = 0.672 vs 193/381 = 0.507, two-sided Fisher exact test, odds ratio = 2.00, p = 5.68 × 10⁻⁷). Most phase place cells were also rate place cells (524/543 = 0.965). We therefore examined phase place cells with and without rate tuning separately and compared each group with rate place cells without phase place tuning. Phase place cells that were also rate place cells were more often pyramidal cells (360/524 = 0.687 vs 179/310 = 0.577, odds ratio = 1.61, p = 0.0016). Phase place cells without rate tuning were more often interneurons (14/19 = 0.737 vs 131/310 = 0.423, odds ratio = 3.83, p = 0.0087).

The proportion of phase place cells also differed between hippocampal subregions (CA1: 517/885 = 0.584, CA3: 29/64 = 0.453, two-sided Fisher exact test, odds ratio = 1.70, p = 0.0490). Dentate gyrus was excluded from the regional comparison because only seven phase-testable neurons were available. Taken together, phase place cells were more often pyramidal cells and their proportion may differ between hippocampal subregions. Phase place cells without rate tuning, however, were mostly interneurons.

**Supplemental Figure 1.**
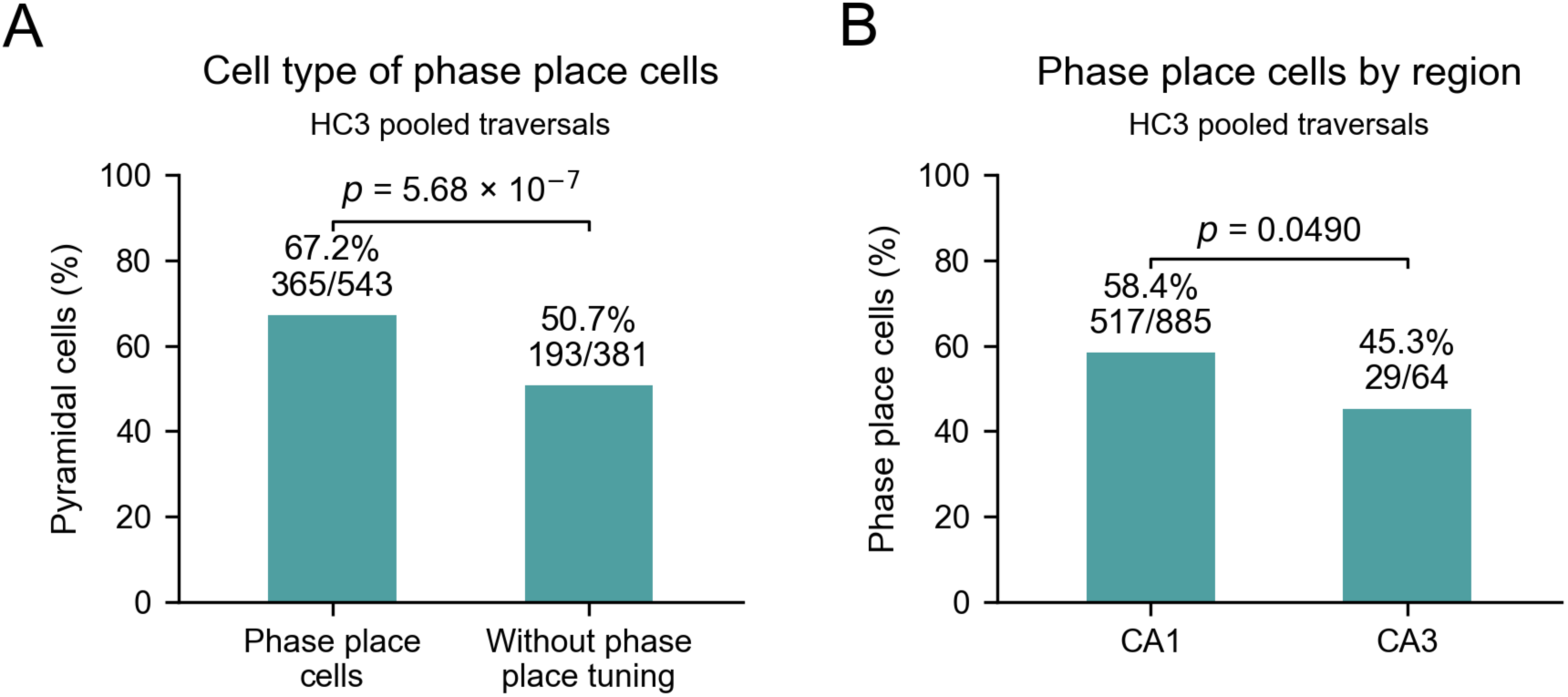
Cell type and subregion of HC3 phase place cells. Phase place classifications are shown for the pooled-traversal HC3 analysis before rate place exclusion. **(A)** Phase place cells were more often putative pyramidal cells than neurons without phase place tuning (365/543 = 0.672 vs 193/381 = 0.507, two-sided Fisher exact test, *p* = 5.68 × 10⁻⁷). Thirty-two phase-testable neurons without a pyramidal/interneuron classification were excluded from this comparison. **(B)** Phase place cells constituted 517/885 CA1 neurons (58.4%) and 29/64 CA3 neurons (45.3%, two-sided Fisher exact test, *p* = 0.0490). Dentate gyrus was excluded from the regional comparison because only seven phase-testable neurons were available.

### Visual concept-task neurons show phase tuning to individual concepts

In the third dataset, we tested whether the same phase-coding idea extended beyond spatial variables. We asked whether neurons in a human visual tuning task showed image-specific phase structure relative to each neuron’s own pre-image baseline. For each neuron, post-image spikes from 0.3 to 1.0 s were compared with a pre-image baseline pooled across images from −0.5 to 0 s. This approach is adapted from methods used to identify rate concept neurons. The retained branch used 5000 permutations and required each tested image to have enough spikes, spikes on at least three (i.e., half) presentations, and no single presentation contributing more than half of the spikes.

The visual tuning task contained putative phase concept neurons under this pre/post baseline analysis. In the inclusive analysis, 11 of 115 eligible neurons showed phase tuning to at least one individual concept (group p=0.0134). Removing benchmark firing-rate tuned images from the candidate image set for the same neuron retained 10 of 114 phase concept neurons (group p=0.0312). This sensitivity removes rate-tuned images, not entire benchmark concept neuron units, so it supports but does not prove a fully rate-independent phase concept population. Thus, CN extends the phase-tuning result from spatial coding to semantic representation in a visual tuning task, while remaining closely tied to firing-rate concept coding (Figure 5).

**Figure 5.**
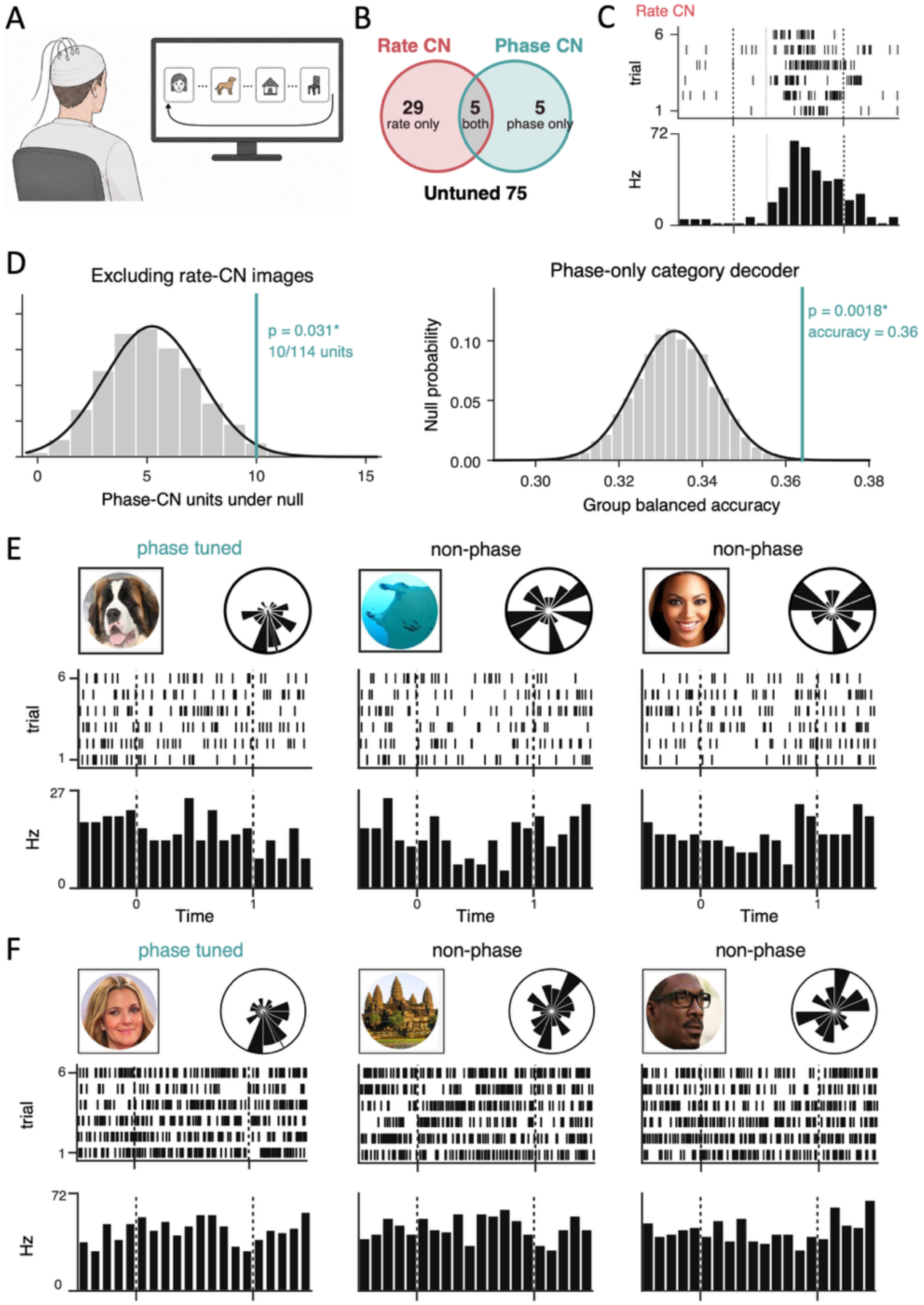
Theta phase carries concept-specific information in the human visual tuning task. **(A)** Participants viewed images of animals, faces, and places while hippocampal single-neuron activity and local field potentials were recorded. **(B)** The Venn diagram shows the overlap between benchmark firing-rate concept neurons (Rate CN) and phase concept neurons (Phase CN) among phase-testable neurons: 29 neurons were rate concept only, 5 showed both forms of tuning, 5 were phase concept only, and 75 showed neither. Phase concept classification was based on the analysis excluding any image that showed firing-rate tuning in the same neuron. **(C)** Example benchmark firing-rate concept response. Raster plots show spikes across the six presentations of an image, and the peri-stimulus histogram shows the corresponding firing-rate response. Dashed vertical lines indicate image onset and offset. **(D)** Left, after removing firing-rate-tuned images from each neuron’s phase concept analysis, 10 of 114 eligible neurons showed significant phase tuning to at least one individual concept (*p* = 0.031). Right, phase features also decoded broad stimulus category (animal, face, or place) above the shuffled-label null (mean balanced accuracy = 36.4%; *p* = 0.0018). Grey histograms show permutation-null distributions and teal lines indicate the observed values. **(E)** Example responses from one neuron. Circular histograms show spike-phase distributions for three images; the image classified as phase tuned is labelled in teal, whereas the two comparison images did not meet the phase-tuning criterion. Raster plots show spikes across the six image presentations, and histograms below show firing rate over time. Dashed vertical lines indicate image onset and offset. **(F)** Same as (E) for a second example neuron.

We next asked whether phase tuning extended from spatial variables to individual visual concepts. In the main analysis, images that already showed firing-rate tuning for a given neuron were excluded from that neuron’s phase concept test. Even after this exclusion, 10 of 114 eligible neurons showed significant phase tuning to at least one individual concept, exceeding the group-level null (*p* = 0.0312). Five of these 10 neurons were also benchmark firing-rate concept neurons through their responses to other images, and five were phase concept neurons only (Fig. 5B). When all images were retained, including those with firing-rate tuning, 11 of 115 eligible neurons were classified as phase concept neurons (*p* = 0.0134). Because the main analysis excluded rate-tuned images rather than entire firing-rate concept neurons, it shows that concept-specific phase tuning can persist for images without detectable rate tuning in the same neuron. These results extend the phase-tuning effect beyond spatial coding to concept-specific representations in the visual task (Figure 5).

At the population level, post-image phase features also carried broader semantic category information. After excluding neuron–image pairs that showed benchmark firing-rate tuning, the decoder distinguished animal, face, and place images above the shuffled-label null (mean balanced accuracy = 36.40% vs. 33.34%; p = 0.00180; Fig. 5).

### Summary of Results

Across datasets, theta-relative spike timing carried information about the represented location or concept. In CityBlock, neurons showed location-specific phase tuning during human navigation, including after rate place cells were excluded. At the population level, phase features from neurons without detectable rate place tuning improved spatial decoding beyond firing rates from the full population, showing that the two signals carried partly nonredundant spatial information. HC3 replicated location-specific phase tuning in the rodent hippocampus and showed that this tuning persisted in a smaller residual population after rate place cells were excluded. Moreover, the strongest phase tuning was more often at the arrival end of traversals, close to reward zones where benchmark spatial rate coding is also typically strongest. In this dataset, however, the firing-rate peaks of rate place cells were not enriched there to the same degree as phase tuning. In the visual tuning task, individual concepts produced image-specific phase structure relative to each neuron’s pre-image baseline, and this effect remained after images with detectable firing-rate tuning were removed from the same neuron. Post-image phase features also decoded broad stimulus category above chance. Together, these findings show that theta-relative spike timing carries spatial and semantic information beyond canonical rate-defined responses.

## Discussion

Across humans and rats, hippocampal theta-relative spike timing carried information beyond canonical rate-defined responses. During human navigation, location-specific phase tuning remained significant after rate place cells were removed. More importantly, phase features from these neurons improved location decoding beyond firing rates from the full population, showing that phase and rate carried nonredundant spatial information. The same rate-excluded phase tuning remained significant in the rat hippocampus. In the visual tuning task, concept-specific phase structure persisted after images with detectable firing-rate tuning were removed. Moreover, phase features alone allowed the decoding of semantic categories. This establishes phase tuning of putative concept neurons beyond a rate code. Together, these results show that theta-relative spike timing can reveal task information that is missed when hippocampal responses are characterized by firing rate alone.

Classical phase precession provides the clearest alternative explanation for the spatial result. As an animal crosses a place field, its spikes advance systematically through theta (O’Keefe and Recce, 1993; Skaggs et al., 1996; Huxter et al., 2008; in humans: Qasim et al., 2021). Phase therefore necessarily varies with position within a rate-defined place field, and this temporal structure can add spatial information to the rate code. A location-specific phase test could consequently identify conventional place cells simply because they precess. We therefore removed rate place cells before asking whether phase distinguished locations. Despite this exclusion, location-specific phase tuning remained significant in both human and rat navigation. The semantic analysis provides the corresponding control outside the spatial domain: concept-specific phase structure remained after rate-tuned images were removed from the same neuron. Our findings therefore extend task-specific phase tuning beyond responses already identified by firing rate.

The rate-exclusion control separates our result from canonical phase precession, but the two may share an input-to-phase mechanism. A place field is not a step function: firing increases toward the field peak, consistent with stronger place-related drive. At the same time, theta modulates the probability that a neuron reaches spike threshold within each cycle. Weak drive may cross threshold only near the most excitable phase, whereas stronger drive can evoke spikes earlier or across a broader portion of the cycle. A gradual increase in place-related drive toward the field peak could therefore allow firing in more inhibited periods thereby shifting spike phase, which could contribute to classical precession. In line with this, controlled depolarization of CA1 pyramidal neurons advances first-spike phase relative to a fixed theta-frequency input, demonstrating that graded depolarizing drive can be converted into theta-relative spike timing (McLelland and Paulsen, 2009). We propose that the same input-to-phase transformation can also operate below a conventional rate threshold. Place- or concept-specific input may be too weak to increase average firing, yet still bias the phase of the spikes that occur. In line with this mechanism, local CA1 activation can generate precession (Sloin et al., 2024), and assembly-specific depolarization shifted preferred phase in a network model (Russo et al., 2024). The same framework therefore applies to semantic tuning: strong concept-specific drive increases firing, whereas weaker drive may alter phase without increasing firing. Phase tuning below the rate threshold and phase precession within a place field may therefore arise when the same graded input interacts with theta-dependent excitability. The same drive-to-phase transformation could thus produce distinct observable regimes above and below a rate-defined threshold. Consistent with this account, some phase place cells showed precession-like phase changes around their preferred location without a detectable rate field. Because location-specific phase locking is tested at every location rather than within a rate-defined field, it can reveal spatial information in the spike timing of neurons that have no place field.

In HC3, the location with the strongest phase consistency was enriched at the arrival ends of traversals, including after rate place exclusion, suggesting that the residual phase signal was spatially organized rather than uniformly distributed along the track. Our data do not measure synaptic drive directly, and the present analyses had limited sensitivity to modest continuous shifts in preferred phase. A direct test of the model would quantify tuning strength continuously and ask whether stronger task-related drive predicts systematic changes in theta-relative spike timing before a rate-defined place or concept response emerges.

Our findings also change how neurons without canonical rate labels should be interpreted. Most phase-testable neurons in the human datasets lacked a canonical rate label: 86% during navigation and 70% during semantic tuning. Rate-only analyses therefore leave a large fraction of recorded spiking activity uncharacterized. Importantly, neurons without place fields can still contribute to spatial population codes (Stefanini et al., 2020) and that spike phase adds information beyond spike counts in primate visual and auditory cortex (Montemurro et al., 2008; Kayser et al., 2009). Our results connect these observations by showing that phase reveals spatial information in neurons without a rate place label and improves decoding beyond the full population rate code. Because action potentials and their postsynaptic effects are metabolically costly (Attwell and Laughlin, 2001), encoding additional information through spike timing without a proportional increase in spike count could in principle be energetically efficient. Whether such efficiency conferred an evolutionary advantage remains speculative.

Our estimates of phase coding are deliberately conservative. We tested phase tuning only in neurons and locations with enough spikes for reliable statistics, and we focused on neurons without rate tuning to separate phase coding from rate coding. Rate-tuned neurons also carry phase information, and neurons that fire too sparsely to be tested may still time their spikes to theta. The phase place cells and phase concept neurons identified here therefore mark a lower bound on how much hippocampal spike timing carries task information.

A remaining question is how a phase code could be organized relative to discrete task events. Behaviorally relevant events can reset or realign hippocampal theta to a common phase (Givens, 1996; Tesche and Karhu, 2000; ter Wal et al., 2021). A theta reset realigns the ongoing oscillation, establishing a common phase reference for the subsequent excitability cycle. Combined with the drive-to-phase relationship described above, this suggests a specific computation: after theta is realigned, neurons receiving stronger task-related drive should reach threshold earlier in the aligned cycle than more weakly driven neurons. Their firing order could therefore rank currently active representations by input strength. The common phase supplied by the reset provides the stable event-related reference required to interpret that order. This hypothesis predicts that continuous tuning strength should predict first-spike order in reset-aligned cycles.

A theta reset could therefore make stronger representations fire earlier than weaker ones within the cycle.

For phase information to influence computation, downstream circuits must also be sensitive to when spikes arrive. When local field potential (LFP) rhythms in connected regions are phase coupled, an upstream spike can reach its target at a reproducible point in the target’s excitability cycle. Anatomical projections can generate interareal coherence (Schneider et al., 2021), and downstream fast-spiking interneurons can lock to the sender’s gamma rhythm (Spyropoulos et al., 2024). A spike arriving during a receptive phase may therefore exert more influence than the same spike arriving at another phase. Spatial information in lateral septal spike phase relative to hippocampal theta is consistent with such a downstream readout (Tingley and Buzsáki, 2018). The principle resembles knocking on a door while loud music plays: a phase-tuned neuron places a knock at the point in the beat when it is most likely to be heard, whereas a rate-coded neuron sends more knocks. Interareal phase coupling could therefore increase the downstream impact of each spike without a large rate increase. This account predicts that the behavioral or postsynaptic impact of a spike should depend on its phase at the receiving circuit, not only on its occurrence.

In conclusion, canonical place cells and concept neurons capture the strongest rate responses but not the full information in the recorded population. Our data show that theta phase coding of location- and concept-specific information remains when the corresponding rate response is not detectable. In human navigation, phase from neurons without a rate place label added spatial information beyond firing rates from the full population. This pattern appeared in humans and rats and for both locations and concepts. A parsimonious framework is that graded task-related drive interacts with theta-dependent excitability: a strong drive changes firing rate, whereas a weaker drive can alter when a spike occurs within the theta cycle. This framework links canonical rate responses, phase tuning, and phase precession without treating them as separate coding schemes. Phase-resolved analysis could therefore reveal information that rate-only analyses miss in datasets containing simultaneous spikes and local field potentials.

### Supplementary Results

Neurons with phase tuning but no rate tuning might have had rate tuning that went undetected because they fired few spikes. We therefore compared, in each dataset, their firing rates in the analysed samples with those of neurons with rate tuning but no phase tuning. In CityBlock, phase place cells without rate tuning fired at a median of 1.5 Hz (n = 12) and rate place cells without phase place tuning at 2.5 Hz (n = 19, Mann-Whitney U test, p = 0.82). In HC3, the corresponding medians were 2.1 Hz (n = 5) and 2.8 Hz (n = 179) among pyramidal cells (p = 0.86), and 26.3 Hz (n = 14) and 17.1 Hz (n = 131) among interneurons (p = 0.12). In the visual task, phase concept neurons without rate tuning fired significantly more than rate concept neurons without phase tuning (15.9 Hz, n = 5, vs 3.9 Hz, n = 29, p = 0.044). Thus, in no dataset did neurons with phase tuning but no rate tuning fire significantly less than neurons with rate tuning but no phase tuning. We therefore found no evidence that low firing rates explain why these neurons lacked rate tuning.

**Supplemental Figure 2.**
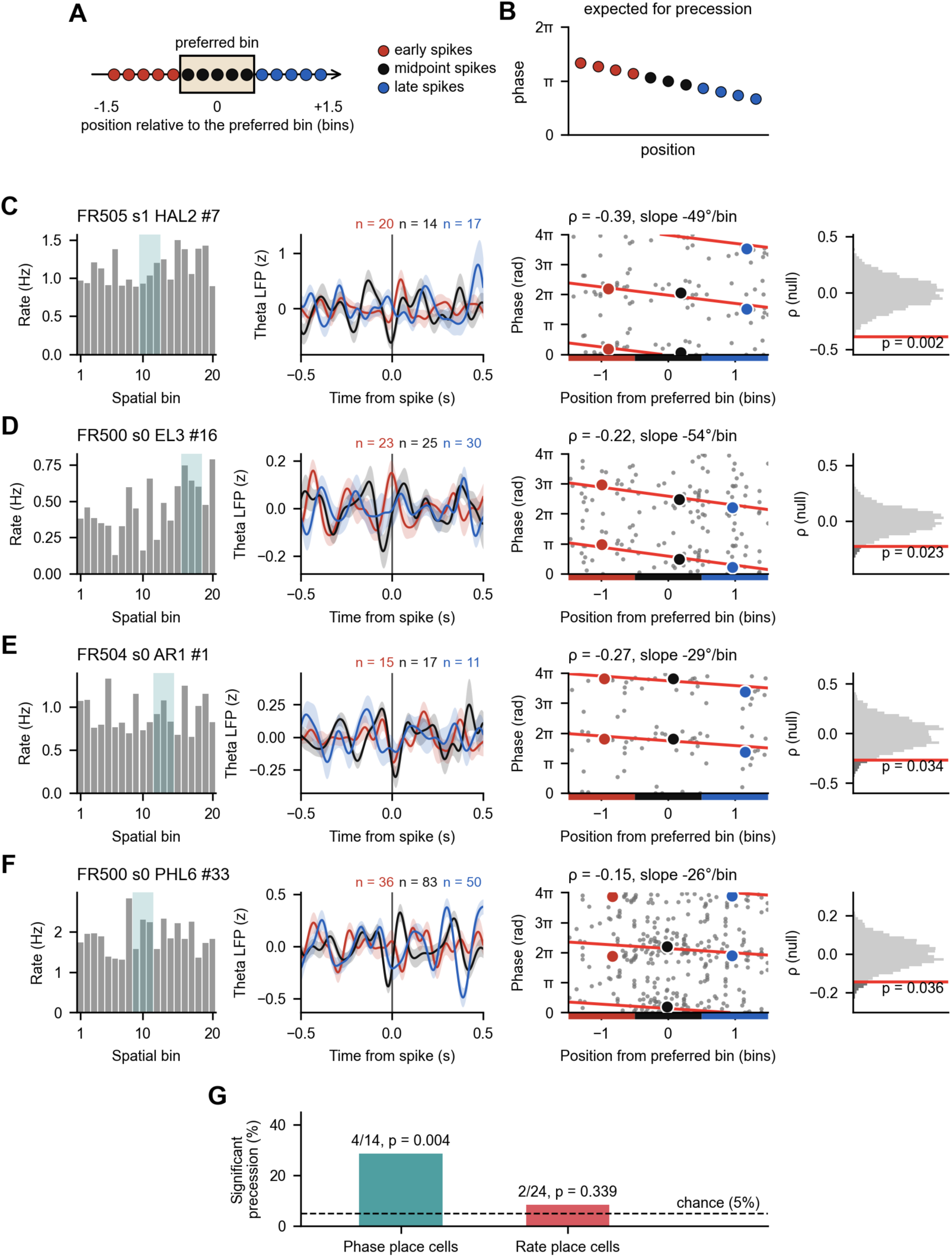
Phase precession in CityBlock phase place cells. (A) Spikes in the analysed window were grouped by position into early spikes (before the preferred bin), midpoint spikes (inside it) and late spikes (after it). (B) With phase precession, spike phase decreases from early to late spikes. (C– F) The four phase place cells with significant phase precession, ordered by p-value. First column, firing rate in each of the 20 spatial bins along the route, with the analysed window shaded. Second column, 3– 12 Hz LFP averaged around early, midpoint and late spikes (z-scored, mean ± SEM), with the number of spikes per group. Third column, theta phase of each retained spike against its position relative to the centre of the preferred bin, plotted over two cycles, with the circular-linear fit (red line, Kempter et al., 2012) and the circular mean phase of each group (coloured dots). Phase 0 corresponds to the theta trough. Fourth column, null distribution of the circular-linear correlation ρ after redrawing spike phases from each neuron’s own phases (5,000 draws), with the observed ρ (red line) and its p-value. (G) Proportion of phase place cells (4/14 = 0.29) and rate place cells (2/24 = 0.083) with significant phase precession, with the 5% chance level (dashed line) and one-sided binomial p-values.

